# Suspensor-derived somatic embryogenesis in Arabidopsis

**DOI:** 10.1101/2020.01.29.924621

**Authors:** Tatyana Radoeva, Catherine Albrecht, Marcel Piepers, Sacco de Vries, Dolf Weijers

## Abstract

In many flowering plants, including *Arabidopsis thaliana*, asymmetric division of the zygote generates apical and basal cells with different fates. The apical cell continues to produce the embryo while the basal cell undergoes a restricted number of anticlinal divisions leading to a suspensor of 6-9 quiescent cells that remain extra-embryonic and eventually senesce. In some genetic backgrounds, or upon ablation of the embryo, suspensor cells can however undergo periclinal cell divisions and eventually form a second, twin seedling. Likewise, embryogenesis can be induced from somatic cells by various genes, but the relation to suspensor-derived embryos is unclear. Here, we addressed the nature of the suspensor to embryo fate transformation, and its genetic triggers. We expressed most known embryogenesis-inducing transcriptional regulators and receptor-like kinases specifically in suspensor cells. Among these, only RKD1 and WUS could induce a heritable twin seedling phenotype. We next analyzed morphology and fate marker expression in embryos in which suspensor division were activated by different triggers to address the developmental paths towards reprogramming. Our results show that reprogramming of Arabidopsis suspensor cells towards embryonic identity is a specific cellular response that is triggered by defined regulators, follows a conserved developmental trajectory and shares similarity to the process of somatic embryogenesis from post-embryonic tissues.

## Introduction

In flowering plants (Angiosperms), embryogenesis is initiated by fertilization of the egg cell. In Arabidopsis, this gives rise to the zygote that undergoes an asymmetric division to form two cells with distinct fate: an apical embryonic cell and a basal extra-embryonic cell from which the suspensor develops. The apical cell then continues to divide in a strictly regular manner to give rise to most tissues and cell types of the seedling plant (Palovaara et al., 2016). Of the approximately 7 suspensor cells, only the uppermost, the hypophysis, contributes to generating the root meristem. The common view is that the suspensor cells supply the growing embryo with nutrients and growth regulators, fix the developing embryo to the micropylar cavity within the seed and may function as a reservoir of embryogenic cells in case the primary embryo fails (Kawashima and Goldberg, 2010; Radoeva and Weijers, 2014). Despite its quiescence under normal conditions, secondary embryos can be formed from suspensor cells in many plant species under specific conditions (Lakshmanan and Ambegaokar, 1984). Suspensor-derived embryogenesis can be experimentally induced by stress treatments (Haccius, 1955) or by impairment of the primary embryo through radiation (Haccius, 1955), mutations (such as *sus* and *twn*; Schwartz et al., 1994; Vernon and Meinke, 1994), genetic ablation (Weijers et al., 2003), by expression of the auxin response inhibitor protein bodenlos (*bdl*; Rademacher et al., 2012) or by laser irradiation (Gooh et al., 2015; Liu et al., 2015). Thus, suspensor cells can be regarded as a dormant pool of stem cells, which can switch to embryo identity in need. Re-initiation of embryonic cell fate in suspensor cells has the advantage that a precise sequence of reprogramming, the possible occurrence of cell autonomy and lateral inhibition as well as stochastic and epigenetic aspects can be analyzed in a predictable fashion in only a few cells of a genetically superior system (Radoeva and Weijers, 2014; de Vries and Weijers, 2017).

The ability of plant cells to be reprogrammed towards embryogenesis has long been recognized and has been the basis for protocols of somatic embryogenesis (Egertsdotter et al., 2019). In the past decades, several factors have been identified that can facilitate or trigger the induction of somatic embryos. Genes like the leucine-rich repeat receptor-like kinase (LRR-RLK) SOMATIC EMBRYOGENESIS RECEPTOR-LIKE KINASE 1 (SERK1) appear to affect the competence to form somatic embryos (Hecht et al., 2001), while transcriptional regulators of the BABY BOOM (BBM) and LEAFY COTYLEDON (LEC) pathway appear to directly induce somatic embryos (Horstman et al., 2017). In addition, genes from the plant-specific RWP-RK domain-containing (RKD) family, involved in maintaining egg cell identity, are able to induce loss of cell identity (Köszegi et al., 2011) or actively promote somatic embryogenesis (Waki et al., 2011) upon ectopic expression. However, these genes were identified and tested in different experimental systems, ranging from Brassica microspores to Arabidopsis meristems and seedlings. This makes it challenging to infer whether these factors are part of the same genetic network or pathway, and it is unclear how these factors, or the process of somatic embryogenesis, relates to the reprogramming of suspensor cells.

Here, we have exploited the simple, predictable suspensor system to address these questions. We have first tested the ability of 16 different genes, representative of the somatic embryo pathways, to induce suspensor-derived twin seedlings. We next compared suspensor-derived embryogenesis induced by three different triggers to define the developmental trajectory underlying reprogramming. We found that a common sequence of events underlies reprogramming. First, suspensor identity is lost, closely connected with activation of cell division. Embryo identity is only gained later, either concomitant with or following the activation of division. Our work shows that suspensor reprogramming is activated by specific triggers, but also reveals a striking similarity between suspensor-derived and other somatic embryogenesis processes.

## Results

### Suspensor embryogenesis requires specific genetic triggers

Several genes have previously been reported to trigger embryogenesis when ectopically overexpressed and have therefore been defined as master embryonic or meristematic regulators (reviewed in Ikeuchi et al., 2013; Radoeva and Weijers, 2014). Nevertheless, their ability to induce embryogenesis has been tested in diverse model systems (Boutilier et al., 2002; Hecht et al., 2001; Waki et al., 2011; Zuo et al., 2002). It is therefore difficult to compare their activities and to address if all indeed trigger embryo identity or some other process contributing to the development of somatic embryos. We decided to use the predictable suspensor-derived embryogenesis as an experimental system to test known embryo inducers for their ability to convert suspensor cells into secondary twin embryos and ultimately into twin seedlings.

We selected sixteen genes to test for their ability to promote embryo initiation. These include transcriptional regulators such as LEC, BBM, AGL15, RKD, WUS and WOX, and the receptor-like kinase SERK1 (Table S1). Each was misexpressed in the developing suspensor using a two-component GAL4/UAS system from the M0171-GAL4 driver line (Rademacher et al., 2012; Radoeva et al., 2016). This same approach previously led to excessive suspensor cell divisions when the *bdl/iaa12*. protein was expressed (Rademacher et al., 2012). However, twin seedlings resulting from ubiquitous *bdl* overexpression were only seen in RPS5A>>*bdl* embryos (Rademacher et al., 2012). While excessively dividing cells in M0171>>bdl suspensors did at times resemble embryo-like structures, twin seedlings were not observed in these embryos (Radoeva et al., 2019).

After transforming transgenes driving each embryogenesis regulator from the GAL4-dependent *UAS* promoter into the M0171 GAL4 driver line, we screened primary transformants for twin seedlings, a phenotype that is not found in wild-type (0%; n>500). This is the most stringent selection criterium for suspensor-derived embryos, given that it not only requires initiation and formation of a second embryo, but also maturation and survival of desiccation, dormancy and germination. Strikingly, while many of these genes had been shown to promote embryogenesis in various conditions, only few could induce twin seedling development in this assay. Only WUS, RKD1 and RKD4 expression led to the recovery of twin seedlings (Fig. 2O; Table S1). The efficiency among these genes was very different: 29% (n=17) of RKD1-expressing transgenics showed twins, in contrast to 3% (n=30) of WUS-expressing and 2% (n=54) of RKD4 expression lines (Table S1). RKD4 resulted in distorted seedlings that did not develop into viable fertile plants (Table S1). The failure to observe twin seedlings with any of the other embryo-promoting genes was not due to the construct used. For example, M0171>>BBM-expressing plants showed morphological distortions at later stage (Table S1), but did not show twin seedlings, twin embryos or periclinal suspensor divisions (data not shown). The RKD1 and WUS-induced twinning was heritable, but phenotypic penetrance was highly variable among RKD1 lines (Table S2).

To determine whether the occurrence of additional, periclinal cell divisions in the suspensor cells coincided with viable twin embryos and seedlings in the same lines we compared phenotype penetrance at different stages. While in 35 % of M0171>>RKD1 embryos (n=366), periclinal suspensor divisions could be observed, only 9% of the viable progeny seedlings were twins (n=1590; Figure 1). Thus, less than one third of the embryos that showed periclinal divisions indeed developed twin embryos. This number is close to what is observed in the *twin1* mutant where 25% of the embryos (n=234) showed periclinal suspensor divisions leading to only 13% of viable seedling twins. The reduced phenotypic penetrance could mean that not all embryo-like structures have embryo identity, but given the delay between suspensor-derived and primary embryo development, it is also possible that spatial constraints or a failure to execute maturation or desiccation programs cause this difference.

**Figure 1:**
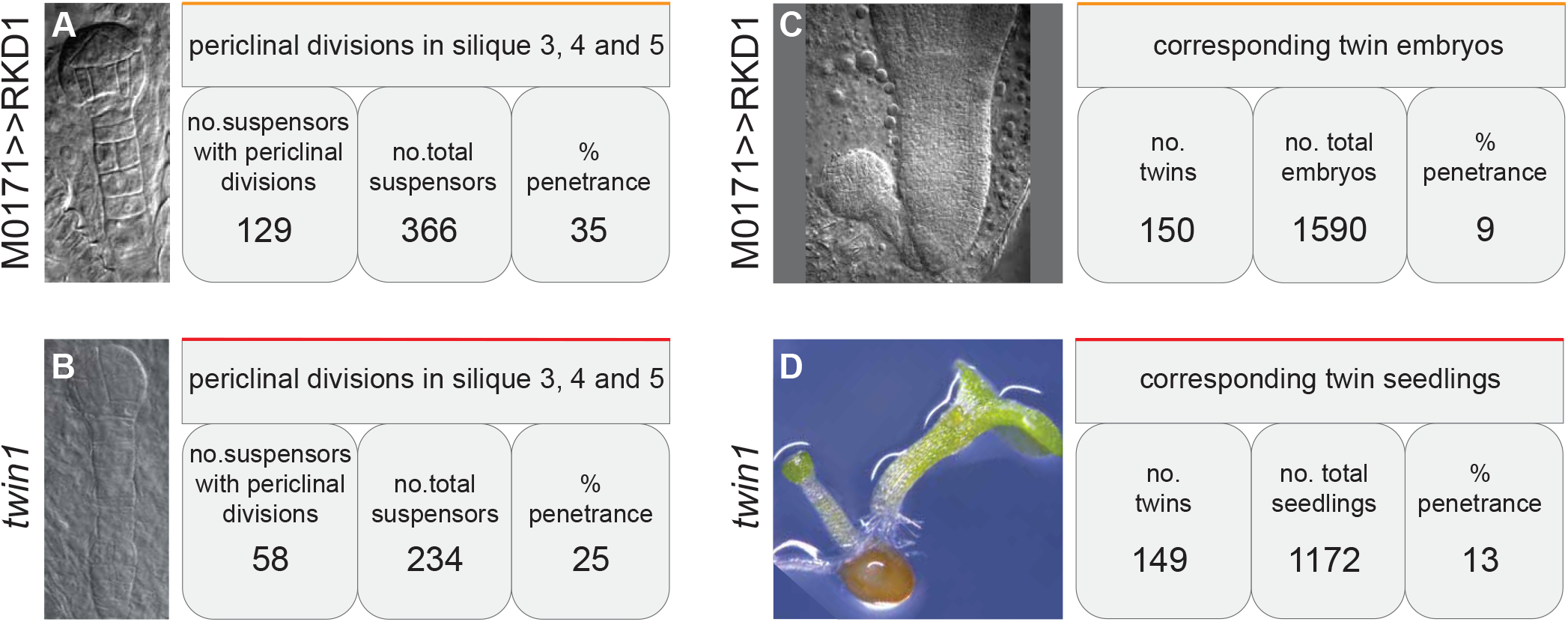
Suspensor division, twin embryo and seedling phenotypes. Frequency of suspensor periclinal divisions compared to twin embryo and seedling formation in M0171>>RKD1 and *twn1* lines.

Thus, at least three transcription factors can induce complete, viable suspensor-derived twin seedlings. Yet, most of the genes tested could not, suggesting that the fate switch in suspensor cells from extra-embryonic to embryonic identity is a specific response that is triggered by a defined set of regulators.

### Diverse cell division patterns can mediate suspensor-derived embryogenesis

Given that multiple independent genotypes each induce suspensor-derived embryos, we asked if the cellular basis for embryo initiation is shared among these genotypes. We therefore compared early embryogenesis in M0171>>RKD1, M0171>>*bdl* and *twin1* genotypes. M0171>>RKD4 was omitted from this analysis because it did not yield a line in which twinning was heritable (Table S1).

In wild type embryos, all suspensor cells are derived from the basal zygote daughter cell through a series of anticlinal divisions (Fig. 2A-D). Only the uppermost hypophysis cell (Fig. 2C) contributes to the root meristem and becomes part of the seedling (Fig. 2E). In all three transgenic (M0171>>*bdl*; M0171>>RKD1) and mutant (*twin1*) genotypes analyzed, the quiescence of the suspensor was disrupted, as expressed by excessive divisions. In M0171>>*bdl* embryos, excessive divisions were found to occur in anticlinal (“normal”), as well as periclinal and oblique planes. While additional anticlinal divisions created longer suspensors, extra periclinal divisions led to the formation of clusters of small cells (Fig. 2I). As described previously (Rademacher et al., 2012), the first periclinal suspensor cell divisions usually occurred at the early globular stage (Fig. 2F). Division defects were however also observed in the pro-embryo (Radoeva et al., 2019). No twin seedlings were observed under standard growth conditions in M0171>>*bdl* lines (Fig. 2J).

**Figure 2:**
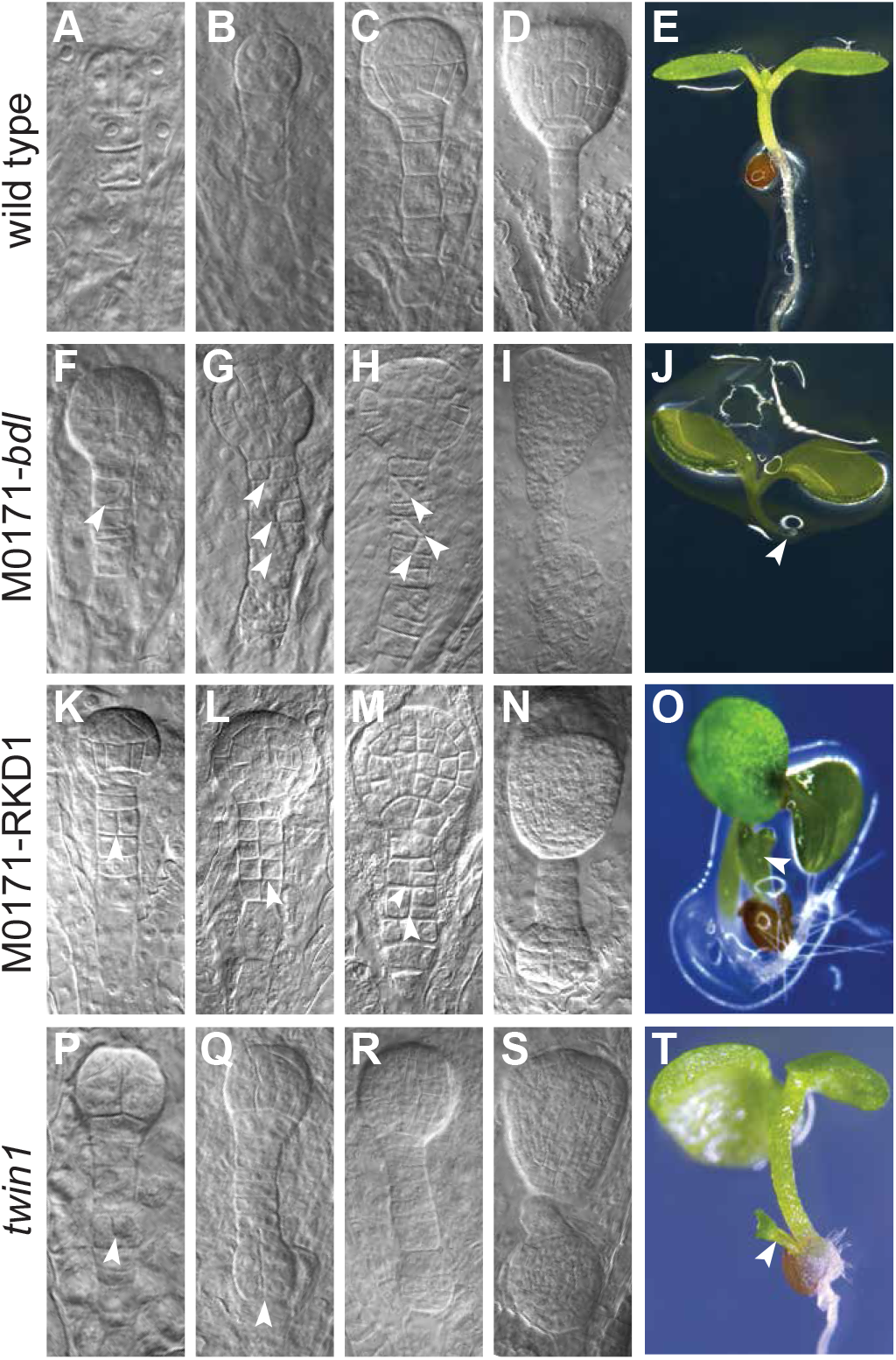
Twin embryo and seedling development. Wild type embryo (A-D) and seedling (E) phenotype, M0171>>*bdl* embryo (F-I) and seedling (J) phenotype, M0171>>RKD1 embryo (K-N) and seedling (O) phenotype, *twn1* embryo (P-S) and seedling (T) phenotype. Embryo images were made from cleared ovules, seedling images by light microscopy. The embryo images in F, K and P were taken at the moment the first periclinal division (arrowhead) was detected. Scale bar represents 10 μ in all panels. Arrowheads point to periclinal divisions, arrowhead in J points to root mutant phenotype, arrows in O and T to secondary twin embryo.

In contrast to the seemingly pleiotropic effect of *bdl* misexpression, M0171>>RKD1 embryos followed a more regular division pattern. Excess divisions in suspensor cells were primarily periclinal, generating ordered multi-layered suspensors, followed by the appearance of embryo-like cell clusters later during development (Fig. 2K-M). While the timing of periclinal suspensor divisions matched that observed in M0171>>*bdl* embryos, no conspicuous defects in the M0171>>RKD1 pro-embryo was detected (Fig. 2N).

The recessive *twin1* mutant showed excessive divisions in the suspensor, and these included both anticlinal (longer suspensors) and periclinal divisions (Fig. 2P). The *twin1* mutant pro-embryo also showed division defects (Fig. 2Q). Embryo-like structures developed in *twin1* suspensors later during development (Fig. 2R), where orientation could be the same as the original embryo, or opposite (Fig. 2S and S5C), and multiple embryos could initiate from the suspensor in a seemingly independent manner (Fig. S5).

We next asked if the ontogeny of cell proliferation in suspensors of these three genotypes were similar. Therefore, we analyzed which suspensor cell showed the first periclinal division, as a clear sign for extra divisions. This analysis showed that first defects occurred more frequently in the top half of the suspensor in M0171>>*bdl* and M0171>>RKD1 lines, while there was no clear preferential origin of the defect in *twin1* embryos (Fig. 3). The hypophysis was excluded from this analysis because *bdl* misexpression specifically interferes with auxin-dependent root formation in this cell (Weijers et al., 2006).

**Figure 3:**
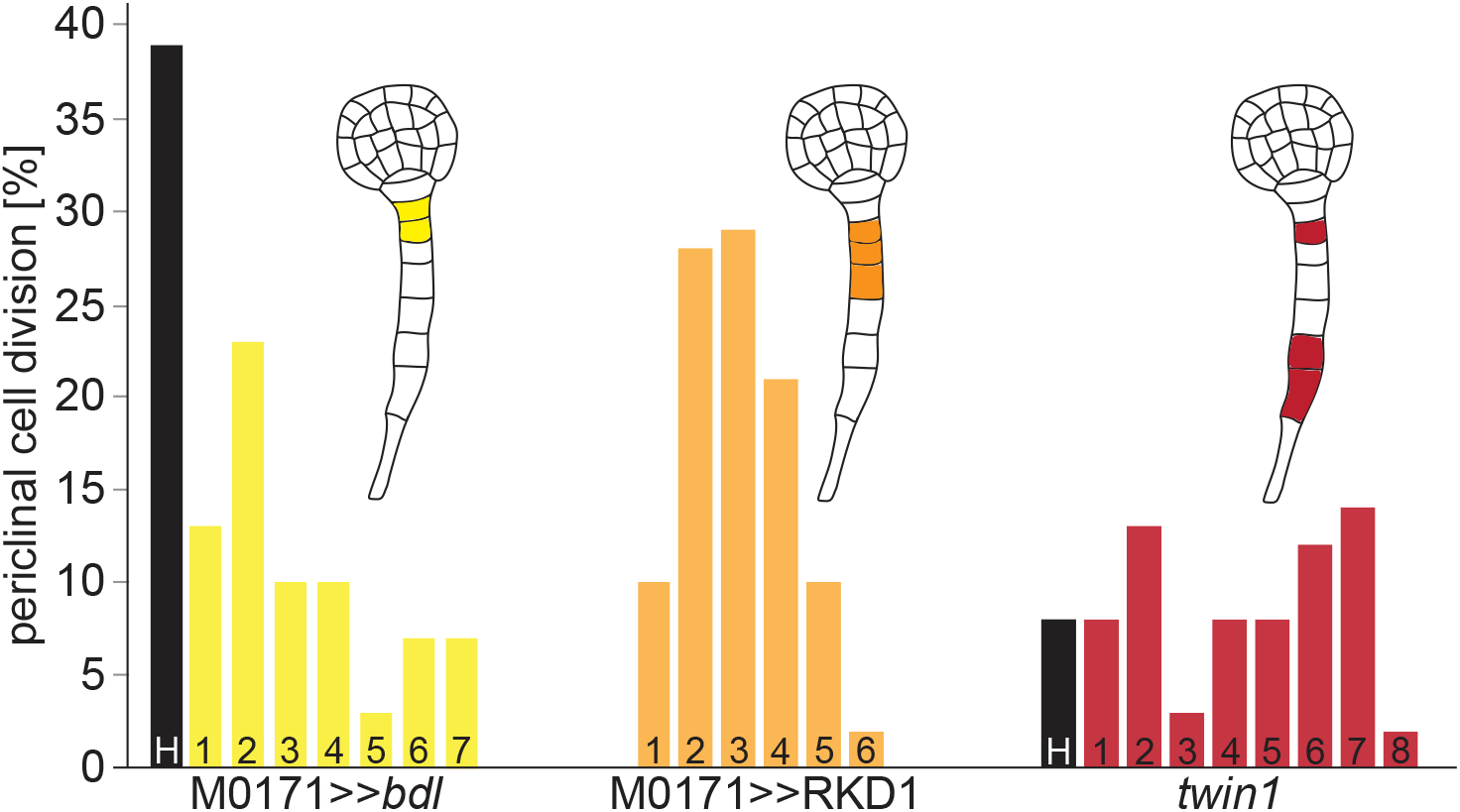
Distribution of periclinal divisions in suspensor cells. Bar diagram of the number of first periclinal divisions observed in M0171>>*bdl*, M0171>>RKD1 and *twn1* suspensors. Bar marked H represents the hypophyseal cell. No periclinal divisions were seen in a comparable number of wild-type embryos.

Based on this phenotypic characterization, all three genotypes that induced suspensor-derived embryos appear to differ with respect to the position of origin in the suspensor, orientation of excessive cell divisions, development of the original pro-embryo and the viability of embryo-like structures. It therefore appears that multiple paths can lead to suspensor-derived embryogenesis.

### Loss of suspensor identity during suspensor-derived embryogenesis

Suspensor-derived embryogenesis is associated with the activation of cell division in suspensor cells, a property shared by all three genotypes tested here. However, it is unclear if the activation of cell division in the suspensor is intimately linked to reprogramming of identities. Alternatively, embryo identity may be activated at any moment after a number of cells have been generated. To address this question, we introduced markers for suspensor or embryo identity into each genotype and analyzed their expression during suspensor-derived embryogenesis.

We first generated a set of markers based on prior publications or on transcriptome data, and evaluated their usefulness as markers of either cell type in wild-type (Table S3). Three markers - pSUC3, pATPase and pWRKY2 – faithfully marked suspensor cells and were introduced in all backgrounds (Fig. S1 and S2). Expression of pATPase∷Venus is strong in suspensor cells at least from the 4-cell stage onwards (Fig. 4A) and is retained during the globular and late heart stage (Fig. 4B,C). In M0171>>*bdl* embryos, expression in suspensor cells is often lost when cells divide periclinally (Fig. 4D). However, not all cells that follow from periclinal division immediately lose pATPase∷Venus expression. This result suggests that, while divisions are accompanied by loss of this suspensor marker, there is likely not to be immediate downregulation following division, or at least not preceding division. In M0171>>RKD1 embryos, pATPase∷Venus expression is reduced throughout the suspensor, and very few cells express the marker at the level found in wild-type suspensors (Fig. 4G). Some residual expression of this marker is retained in suspensor cells, even when these divide periclinally (Fig. 4H). Likewise, in *twin1* suspensors, the pATPase marker is also mostly lost when cells form embryo-like structures (Fig. 4K). Analysis of pSUC3 and pWRKY2 markers in M0171>>*bdl* and M0171>>RKD1 embryos showed comparable results (Fig. S2). Hence, the activation of cell divisions in all three genotypes are indeed associated with loss of suspensor markers. However, there is no immediate shutdown of marker expression upon the first periclinal division. Given that the suspensor cell cycle is approximately 15 h (Gooh et al., 2015), while the half-life of the Venus variant used here is estimated to be about 24 h (Snapp, 2009), it is well possible that Venus signal is retained in divided cells, even if there is no more transcription after division. To address this question, we quantified Venus signals in periclinally divided suspensor cells, and found these to be about half of that in non-dividing and anticlinally divided cells (Fig. S3). This suggests that expression of suspensor-specific promoters is switched off during or after periclinal division.

**Figure 4:**
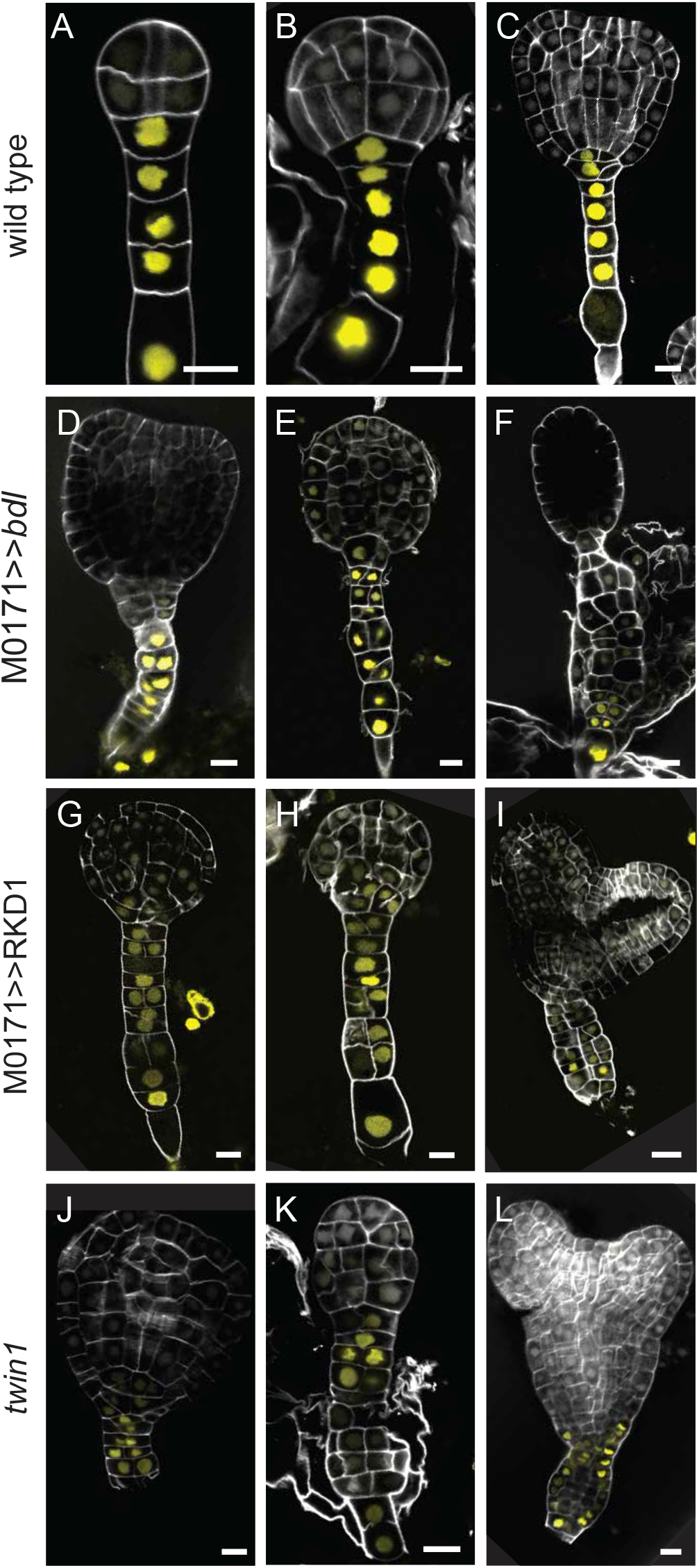
Loss of suspensor marker expression after periclinal suspensor cell divisions. Expression of the pATPase∷Venus suspensor marker in wild-type (A-C), M0171>>*bdl* (D-F), M0171>>RKD1 (G-I) and *twn1* embryos (J-L). All embryos were released from developing seeds and imaged by confocal microscopy. Scale bar represents 10 μm in all panels.

### Division likely precedes loss of suspensor identity

Given that periclinal division is associated with loss of suspensor marker gene expression, and with the initiation of embryo-like structures, an important question is whether the divisions are a consequence or a cause of reprogramming towards embryogenesis. In the former scenario, one would expect loss of suspensor markers before cells first divide periclinally. Since this would be difficult to infer from observing multiple embryos due to the variation of fluorescence levels within and between embryos, we used a live imaging approach to establish timing of division and expression of the pATPase marker in *twin1* mutant embryos. In wild-type, embryos, occasional anticlinal divisions were observed during the observation time of 64 h (Fig. 5A), consistent with prior analysis of divisions in wild-type embryos (Gooh et al., 2015). Levels of pATPase marker fluorescence did not change noticeable either before, during or after these anticlinal divisions (Fig. 5A). We observed several periclinal divisions in *twn1* mutant embryos (Fig. 5B, C). In these cells, we did not however detect a change in pATPase expression prior to the periclinal division. Rather, expression decreased in daughter cells after the division. In some cases, expression of the marker was re-activated in one of the daughter cells (Fig. 5C). In conclusion, the loss of suspensor marker expression occurs after, not before or during, the periclinal division, which suggests that divisions are not the consequence of reprogramming. Rather, these divisions provide the cells in which reprogramming can occur.

**Figure 5:**
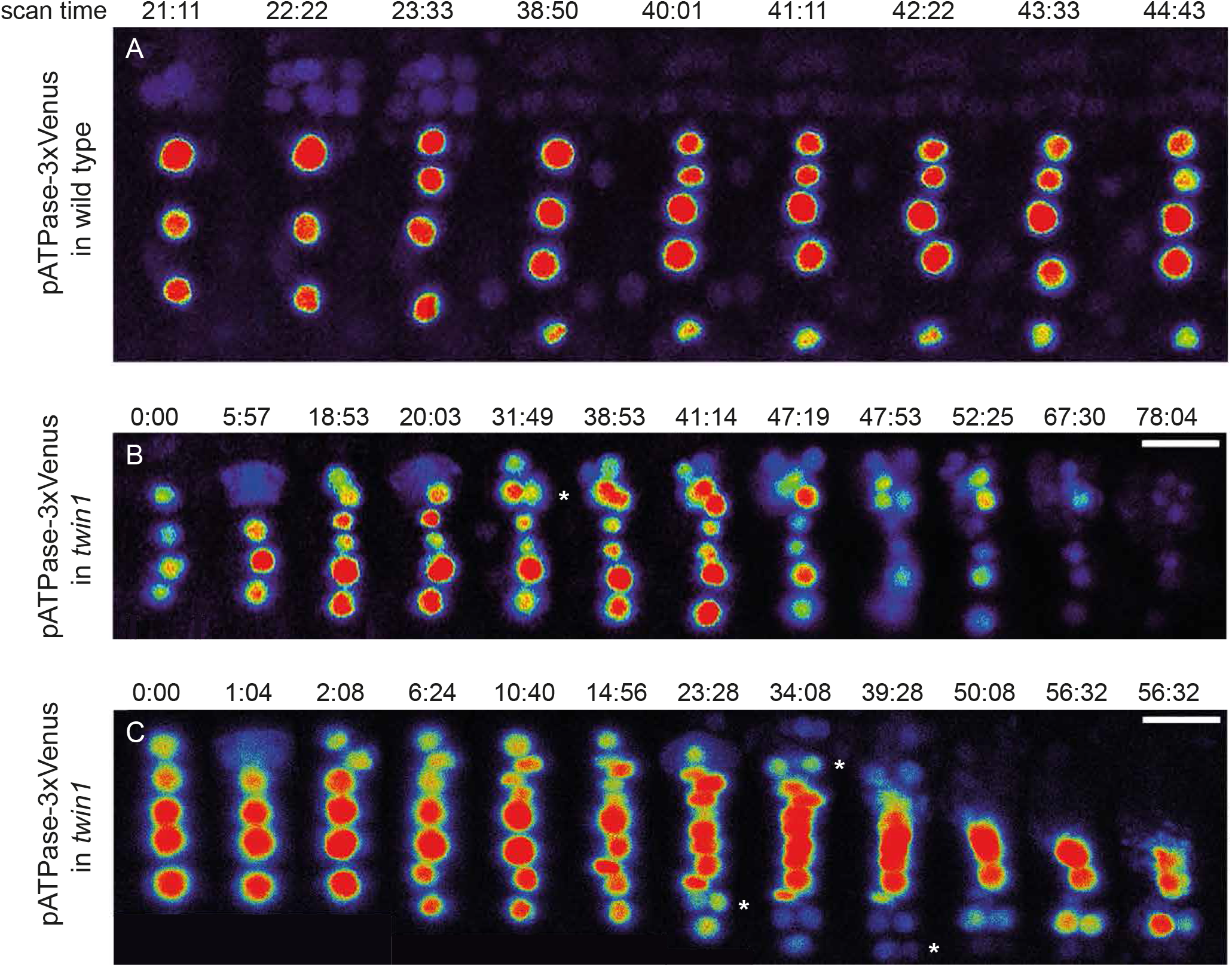
Live imaging of suspensor marker expression in *twn1*. Time-lapse recordings of pATPase∷Venus expression in wild-type (A), and in two representatives of *twn1* (B,C). Up to approximately 45-50 h of recording no loss of pATPase∷Venus signal intensity was observed in wild type embryos.

### Activation of embryo identity in suspensor-derived embryos

To determine when newly divided cells in the suspensor switch on an embryo program, we analyzed the expression of the pDRN∷Venus marker in the three genotypes. pDRN∷Venus was selected from a larger collection (Table S3) based on its specificity and early expression in the wild-type pro-embryo at the 4-cell stage (Fig. 6A). Following its activation in the apical cell(s), DRN expression persists in the apical half of the early globular embryo (Fig. 6B) and becomes restricted to the shoot apical meristem (Fig. 6C). Despite clear expression in the pro-embryo, we could not detect activation of the DRN marker in dividing cells (Fig. 6D) and proliferating cell clusters (Fig. 6E, F) of M0171>>*bdl* suspensors. It should be noted though, that DRN is a direct target of the auxin response factor MP (Cole et al., 2009), whose expression is activated in proliferating suspensor cells (Rademacher et al., 2012). It is therefore likely that *bdl* expression in the suspensor will inhibit DRN expression irrespective of whether cells acquire embryo identity.

**Figure 6:**
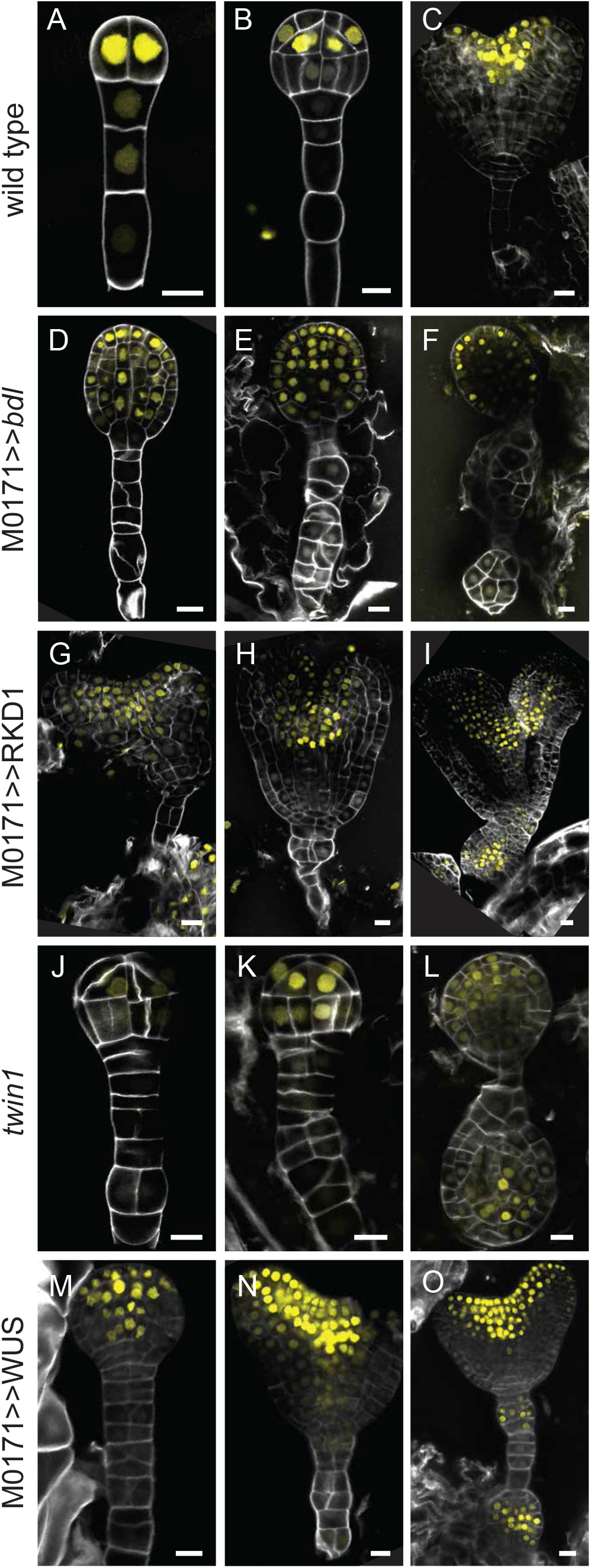
Activation of embryo marker expression in suspensor-derived embryos. Expression of the pDRN∷Venus pro-embryo marker in wild-type (A-C), M0171>>*bdl* (D-F), M0171>>RKD1 (G-I) and *twn1* embryos (J-L). All embryos were released from developing seeds and imaged by confocal microscopy. Scale bar represents 10 μm in all panels.

In contrast, periclinal divisions in suspensors of M0171>>RKD1 embryos were accompanied by activation of DRN expression (Fig. 6G, H). In most embryos DRN expression was not seen until a small cluster of proliferating cells had been established in the suspensor (Fig. 6I). The same was observed in *twin1* embryos (Fig. 6J-L), and was confirmed in M0171>>WUS embryos (Fig. 6M-O). To address whether the cells observed after the loss of suspensor identity and before acquisition of embryo identity followed a pathway mimicking egg cell identity, expression of a reproductive expression cassette-FGR7.0 (egg cell, synergids and central cell; (Völz et al., 2013) was crossed with the *twin1* line. No expression of any of the markers could be detected during periclinal divisions and the formation of twin embryos (Fig. S4.)

The analysis of suspensor and embryo markers in four genotypes reveals that the process of reprogramming suspensor cells towards embryo identity is marked by periclinal cell divisions, loss of suspensor markers and gain of an embryo marker. It appears that in most cases, cell divisions and loss of suspensor identity occurs well before an embryo marker is activated. This could of course be caused by difficulties in detecting early DRN expression due to low phenotype penetrance and low expression levels. However, these observations are also consistent with a scenario where reprogramming involves three distinct processes: loss of suspensor identity, cell proliferation and gain of embryo identity.

### Direct conversion of suspensor into embryo identity?

The analysis of the DRN marker in M0171>>RKD1, M0171>>WUS and *twn1* embryos show that embryo identity is activated in newly formed cell clusters, but it is difficult to define the timing of activation relative to divisions. This is mainly due to the limited phenotypic penetrance, which complicates detection of the earliest events. In propagating primary and secondary embryos from these three genotypes, we recovered lines that show a strongly increased phenotypic penetrance that allow to address this question.

Since the suspensor-derived (secondary) embryo is initiated when the original (primary) pro-embryo is at globular or heart stage (Fig. 2), it is delayed and therefore smaller than the primary embryo at maturity. This causes a size difference between the larger primary and the smaller secondary seedling in *twin1*, M0171>>RKD1 and M0171>>WUS lines (Fig. 7A). We separately propagate primary and secondary seedlings from these genotypes and tested their progeny for the penetrance of twin phenotypes. Strikingly, the progeny from secondary seedlings showed much higher phenotype penetrance compared to the primary seedling-derived progeny in both M0171>>RKD1 and M0171>>WUS lines (Fig. 7A). In the secondary seedling-derived lines, triplet suspensor-derived embryos were occasionally seen, in rare instance giving rise to triplet seedlings (Fig. S5). In contrast, this difference was not observed in the *twn1* mutant (Fig. 7A), suggesting an epigenetic component that acts on the regulation of the M0171 GAL4 driver. Nonetheless, we leveraged the increased phenotypic penetrance in secondary seedling-derived M0171>>RKD1 and M0171>>WUS embryos to help identify the earliest stages of activation of the DRN marker. Indeed, we more readily identified periclinal suspensor cell divisions and substantially earlier expression of the DRN∷Venus marker. This could be observed as early as after the first periclinal division in both daughter cells (Fig. 7B, C, E, F). We re-examined the *twin1* mutant in light of this observation, which revealed that early DRN∷Venus expression also occurs in this mutant (Fig. 7D). Thus, in addition to the ‘late’ DRN expression in cell clusters, there appears to also be a more direct conversion into embryo identity.

**Figure 7.**
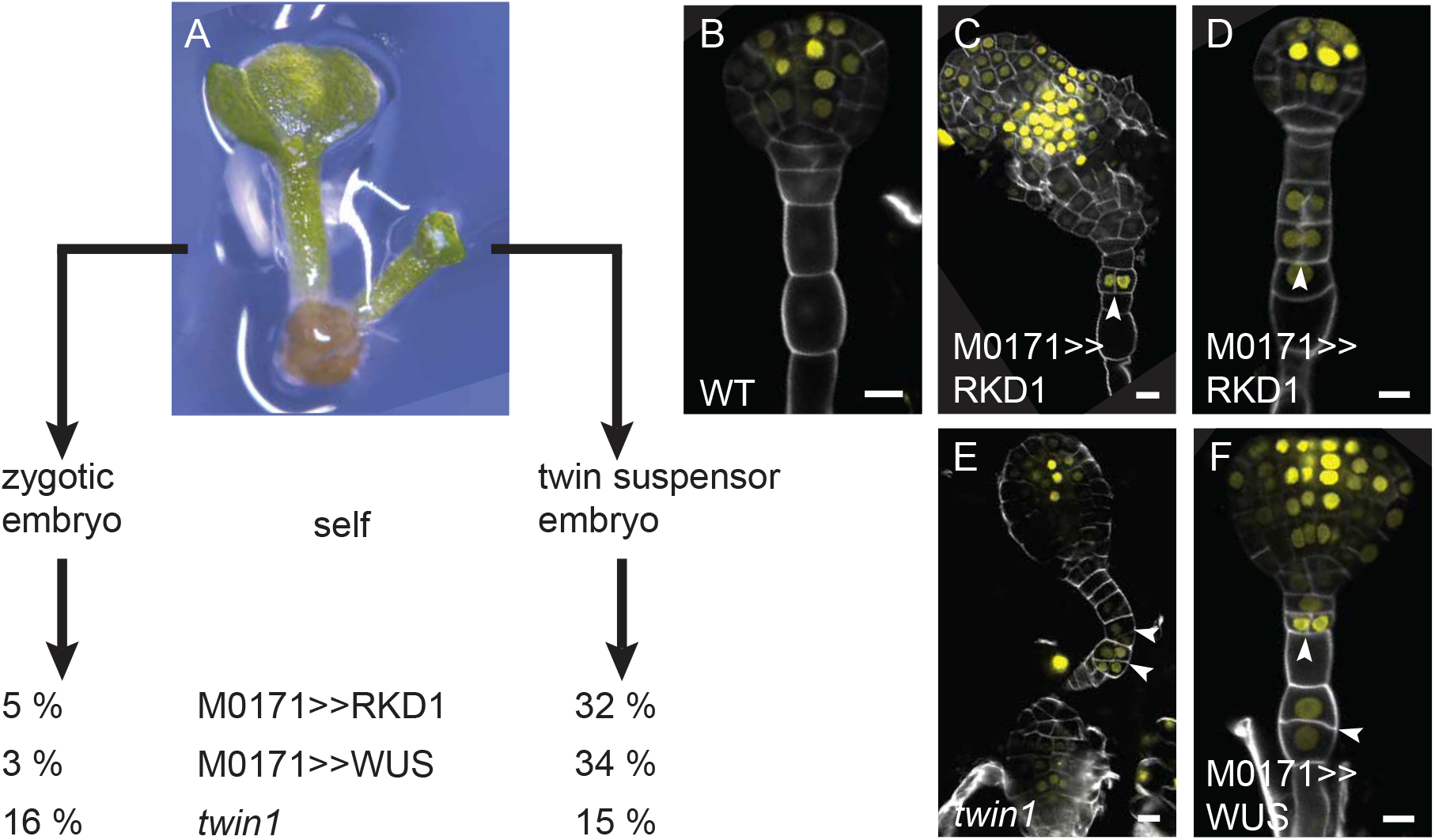
Early embryo fate conversion in high-penetrance lines. Three rounds of selfing using *twin1*, M0171>>RKD1 and M0171>>WUS, expressing the pDRN∷Venus marker produced a collection of lines that derived from the zygotic or the suspensor-derived twin embryo. Between 7 and 16 lines (over 1300 embryos in total) per transgene were analysed for the indicated penetrance of the twin embryo phenotype (A). Early pDRN∷Venus expression was recorded in wt embryos (B), the third generation of twin suspensor embryo derived lines of M0171>>RKD1 (C, D) *twin1*(E) and M0171>>WUS (F, G). All embryos were released from developing seeds and imaged by confocal microscopy. Scale bar represents 10 μm in all panels. Arrowheads indicate single suspensor cells that express the pro-embryo marker pDRN∷Venus.

## Discussion

The occurrence of twin seedlings is a rare property in Arabidopsis, previously found only in the recessive *twin1* and *twin2* mutants (Vernon and Meinke, 1994) and upon inhibition of auxin response (Rademacher et al., 2012). Here, we explored a candidate gene approach employing suspensor-specific expression of genes known to promote somatic embryogenesis. This revealed that three genes, RKD1, RKD4 and WUS were able to induce twin seedlings. Of these, WUS and the egg-cell expressed gene RKD1 resulted in a heritable twin embryo and seedling phenotype.

One of the surprising findings is that transcription factors known to maintain embryo identity such as BBM (Boutilier et al., 2002), AGL15 (Harding et al., 2003), LEC (Braybrook and Harada, 2008) and members of the WOX family (Haecker et al., 2004) did not induce twins. Apart from possible transgene silencing effects, a plausible interpretation is that the activity of embryo-inducing genes is far more dependent on cellular context than anticipated. Context-dependent action has been described for genes belonging to the BBM-AGL15-LEC pathway, that appear to be more active in immature zygotic embryos than in mature seedlings (Horstman et al., 2017). Apparently, context-dependence also extends in the opposite direction to much earlier stages of embryo development as analyzed here.

The WUSCHEL gene is a homeobox-containing transcription factor that maintains the undifferentiated state of stem cells in the shoot apical meristem (Laux et al., 1996; Mayer et al., 1998). Later it was found in an activation tagging screen as an effective inducer of somatic embryos from seedling roots (Zuo et al., 2002). It is therefore remarkable that suspensor-enhanced expression of a gene promoting the undifferentiated state results in countering the normally imposed block of embryogenic potential of the suspensor cells. Remarkably, in our screen WUS is the only gene reported to promote embryogenesis out of context in root cells and also in suspensor cells. In a genome-wide analysis of genes expressed in Arabidopsis somatic embryos, compared to leaf tissue and undifferentiated callus cells, WUS was found to be upregulated in somatic embryos (Wickramasuriya and Dunwell, 2015). Therefore, it appears that the cellular states underlying meristem pluripotency and embryogenesis share a common trigger.

RKD1 is a member of a small Arabidopsis gene family of RWP-RK domain-containing proteins with transcription factor activity that were originally found as genes preferentially expressed in wheat egg cells (Köszegi et al., 2011). Ectopic expression of RKD1 resulted in callus formation with egg cell characteristics. Extensive analysis of multiple mutant combinations did not provide a clear role for RKD1 in female gametogenesis (Tedeschi et al., 2017) and no evidence was provided for RKD1 functions beyond potentially maintaining egg cell identity. Loss-of-function alleles of another member of this family, RKD4, impairs zygote cell elongation and subsequent early divisions. Ectopic expression of RKD4 induces callus from which somatic embryos can form after depleting RKD4 (Waki et al., 2011). These results suggest a more general role of RKD proteins in gametophyte identity and early embryogenesis (Koi et al., 2016). We found that transient RKD1 expression in suspensor cells leads to heritable twin seedling formation. RKD4 had similar but more limited potential, as the single transgenic twin line did not show heritability of the phenotype. Given the proposed role of RKD1 in promoting egg cell identity, a plausible possibility would be that RKD1 expression caused suspensor cells to revert back to an egg cell state. This was not directly tested, but an egg cell marker was not activated during reprogramming in the *twn1* mutant. It is therefore likely that RKD1 expression does not simply trigger egg cell identity in suspensor cells., and that its activity is also context-dependent.

It is intriguing that the only two genes that could efficiently convert suspensor cells into embryogenic are those that appear to have a role in promoting an undifferentiated state in either the shoot meristem cells or in the egg cell.

A key event in induction of somatic embryos in plant tissue culture is an event that has long been considered dedifferentiation. Since it is unlikely that cells entirely loose all aspects of their original identity, this event is perhaps better viewed as reprogramming. What follows is a mass of rapidly dividing cells (Fehér, 2019). Such cells exhibit a callus-like transcriptome (Che et al., 2006; Xu et al., 2012) and transcription factors such as WIND (Iwase et al., 2011) have been identified that promote subsequent steps in regeneration (Iwase et al., 2015). Few studies have addressed the question whether reprogramming followed by the acquisition of a new cell fate such as ‘embryogenic’ first requires erasure of the previous somatic cell fate. Our results show that upon initiating periclinal cell division in suspensors, suspensor marker gene activity was generally reduced or totally absent. In the context of suspensor reprogramming, loss of existing cell identity is therefore indeed the first sign of cellular fate change. Propagating suspensor-derived RKD1 or WUS seedlings to later generations resulted in lines that showed increased penetrance of the twin seedling phenotype. Clearly, this suggests the existence of an epigenetic component involved in the fate conversion of suspensor cells into embryogenic cells. It should be noted that this effect was observed in M0171>>RKD1 and M0171>>WUS lines, but not *twn1*. The insertion site of the transgene in the M0171 could not be identified (Radoeva et al., 2016), and may reside in a genomic area with repeats or high GC content, perhaps sensitive to epigenetic phenomena. On the other hand, it is also possible that the process of reprogramming itself involves epigenetic components. Indeed, explants derived from somatic embryos often exhibit an increased frequency of embryogenic cell formation compared to original explants (reviewed in Méndez-Hernández et al., 2019). Whether a link exists between this phenomenon and the recently discovered role of chromatin remodeling in embryogenic cell formation (reviewed in De-la-Peña et al., 2015; Guo et al., 2020) remains to be determined. In *twin1* and in the high-penetrance RKD1 and WUS lines, DRN expression was activated almost immediately upon suspensor cell division, suggesting a direct conversion of suspensor cell into embryo fate. What this result shows is that reprogramming can, but must not, involve an intervening period of cell proliferation. Based on the similarities described above, we propose that suspensor-derived embryogenesis is closely related to the classical process of somatic embryogenesis.

## Acknowledgements

The authors would like to thank Rita Gross-Hardt for sharing the reproductive cell marker FGR7.0, David Meinke for the *twin1* mutant, Thijs de Zeeuw for advice on live imaging and Naomi Weertman for experimental support. This work was supported by the Netherlands Organization for Scientific Research (NWO; ALW-NSFC Grant 846.11.001 to D.W.).

## Materials and Methods

### Plant material and growth conditions

The M0171 GAL4/GFP enhancer trap line was generated by Dr. Jim Haseloff in the C24 ecotype (Haseloff, 1999) and was obtained through the Nottingham Arabidopsis Stock Center (NASC). All transcriptional Venus fusion lines and the pUAS-gene fusion lines were generated in Columbia-0 (Col-0) ecotype.

Seeds were surface sterilized in 25% bleach/75% ethanol solution for 10 minutes and were afterwards washed twice with 70% ethanol and once with 100% ethanol. Dried seeds were subsequently plated on half-strength Murashige and Skoog (MS) medium and the appropriate antibiotic (50 mg/l Kanamycin, 15 mg/l Phosphinothricin or 0.1 mg/l Methotrexate) for selection of transgenic seeds. After 24 hours incubation at 4°C, the plants were cultured under long-day (16h light, 8h dark) conditions at 22°C.

### Cloning

All cloning was carried out using the Ligation Independent Cloning (LIC) system and the vectors used are previously described (de Rybel et al., 2011; Wendrich et al., 2015). For generating the pUAS-fusion lines for M0171-drive misexpression, genomic fragments spanning the entire coding sequences were amplified from genomic DNA using Phusion Flash PCR Master Mix (Thermo Scientific) and cloned into vector pPLV132. To generate the transcriptional fusions, up to 3kb fragments upstream of the ATG were amplified from genomic DNA. After sequencing, the constructs were transformed into M0171>>RKD1, UAS-*bdl*, *twin1* and M0171>>WUS lines by floral dipping (de Rybel et al., 2011). All primers used for cloning can be found in Table S4.

### Microscopy and sample preparation

Differential Interference Contrast (DIC) and confocal microscopy were carried out as previously described (Llavata-Peris et al., 2013) with minor modifications. For DIC imaging, ovules were isolated in chloral hydrate solution (chloral hydrate, water and glycerol, 8:3:1 w/v/v). After short incubation, the embryos were observed on a Leica DMR microscope equipped with DIC optics. For confocal imaging, ovules were isolated in 1x phosphate solution saline (PBS) containing 4% paraformaldehyde, 5% glycerol and 0.1% SCRI Renaissance Stain 2200 (R2200; Renaissance Chemicals, UK) for counterstaining of embryos. The embryos were taken out of the ovules by gently pressing the coverslip of slides containing ovules. R2200 and Venus fluorescence were visualized by excitation at 405 nm and 514 nm and detection between 430-470 nm and 524-540 nm, respectively. Confocal imaging was performed on a Leica SP5 II system equipped with Hybrid detectors (HyD).

### Live embryo imaging

For live imaging, the procedures described by (Gooh et al., 2015) were employed with a number of modifications. M0171>>*bdl*, M0171>>RKD1 and *twin1* lines showing a high penetrance of the twin seedling phenotype and that also expressed pATPase∷Venus markers were selected. Approximately 50-80 ovules were isolated and incubated on 300 μ polydimethylsiloxane (PDMS) microcage arrays, modified by cutting a small channel in the devise to allow better exchange with the surrounding Nitsch medium supplemented with 5% w/v trehalose (Gooh et al., 2015). This resulted in ovules remaining alive and growing up to 300 h. Suspensor markers remained visible for at least 110 h of culture time using an hourly schedule of illumination.

Live embryo tracking was done on a Leica SP8 with an inverted table controlled by the LAS AF and LAS X programs. A 20x water objective using 20% glycerol to prevent evaporation during long acquisition times was used. To visualize Venus fluorescence, excitation was done at 514 nm, 20% laser power and acquisition between 535 and 570 nm. After manually pinpointing ovule positions, the program collected 10 z-stack images at 10 μ spacing with the most intense image at the center, recorded every hour and about 20 ovules per microcage. Image data were optimized to obtain z-projections that were mounted in sequence. All projections were evaluated for the occurrence of anticlinal (wild type) and periclinal (mutant or transgene) suspensor cell divisions. Quality of fluorescent images was scored using an ad hoc scaling system between 0 and 4.

## References

Boutilier, K., Offringa, R., Sharma, V.K., Kieft, H., Ouellet, T., Zhang, L., Hattori, J., Liu, C.M., Van Lammeren, A.A.M., Miki, B.L.A. and Custers, J.B., van Lookeren Campagne, M.M. (2002). Ectopic expression of BABY BOOM triggers a conversion from vegetative to embryonic growth. Plant Cell. 14, 1737–1749. doi: 10.1105/tpc.001941.

Braybrook, S.A., and Harada, J.J. (2008). LECs go crazy in embryo development. Trends Plant Sci. 13, 624–630. doi: 10.1016/j.tplants.2008.09.008.

Che, P., Lall, S., Nettleton, D., and Howell, S.H. (2006). Gene expression programs during shoot, root, and callus development in Arabidopsis tissue culture. Plant Physiol. 141, 620–637. doi: 10.1104/pp.106.081240.

Cole, M., Chandler, J., Weijers, D., Jacobs, B., Comelli, P., and Werr, W. (2009). DORNRÖSCHEN is a direct target of the auxin response factor MONOPTEROS in the Arabidopsis embryo. Development. 136, 1643–1651. doi: 10.1242/dev.032177.

De-la-Peña, C., Nic-Can, G.I., Galaz-Ávalos, R.M., Avilez-Montalvo, R., and Loyola-Vargas, V.M. (2015). The role of chromatin modifications in somatic embryogenesis in plants. Front. Plant Sci. 18, 6:635. doi: 10.3389/fpls.2015.00635.

Egertsdotter, U., Ahmad, I., and Clapham, D. (2019). Automation and scale up of somatic embryogenesis for commercial plant production, with emphasis on conifers. Front. Plant Sci. 10, 10. doi: 10.3389/fpls.2019.00109.

Fehér, A. (2019). Callus, dedifferentiation, totipotency, somatic embryogenesis: What these terms mean in the era of molecular plant biology? Front. Plant Sci. 10, 10. doi: 10.3389/fpls.2019.00536.

Gooh, K., Ueda, M., Aruga, K., Park, J., Arata, H., Higashiyama, T., and Kurihara, D. (2015). Live-Cell Imaging and Optical Manipulation of Arabidopsis Early Embryogenesis. Dev. Cell. 34, 242–251. doi: 10.1016/j.devcel.2015.06.008.

Guo, H., Fan, Y., Guo, H., Wu, J., Yu, X., Wei, J., Lian, X., Zhang, L., Gou, Z., Fan, Y., et al. (2020). Somatic embryogenesis critical initiation stage-specific mCHH hypomethylation reveals epigenetic basis underlying embryogenic redifferentiation in cotton. Plant Biotechnol. J. doi: 10.1111/pbi.13336.

Haccius, B. (1955). Experimentally Induced Twinning in Plants. Nature. 176, 355–356. doi: 10.1038/176355a0.

Haecker, A., Groß-Hardt, R., Geiges, B., Sarkar, A., Breuninger, H., Herrmann, M., and Laux, T. (2004). Expression dynamics of WOX genes mark cell fate decisions during early embryonic patterning in Arabidopsis thaliana. Development. 131, 657–668. doi: 10.1242/dev.00963.

Harding, E.W., Tang, W., Nichols, K.W., Fernandez, D.E., and Perry, S.E. (2003). Expression and Maintenance of Embryogenic Potential Is Enhanced through Constitutive Expression of AGAMOUS-Like 15. Plant Physiol. 133, 653–663. doi: 10.1104/pp.103.023499.

Haseloff, J. (1999). GFP variants for multispectral imaging of living cells. Methods Cell Biol. 58, 139–151. doi: 10.1016/s0091-679x(08)61953-6.

Hecht, V., Vielle-Calzada, J.P., Hartog, M. V., Schmidt, E.D.L., Boutilier, K., Grossniklaus, U., and De Vries, S.C. (2001). The arabidopsis SOMATIC EMBRYOGENESIS RECEPTOR KINASE 1 gene is expressed in developing ovules and embryos and enhances embryogenic competence in culture. Plant Physiol. 127, 803–816. doi: 10.1104/pp.010324.

Horstman, A., Li, M., Heidmann, I., Weemen, M., Chen, B., Muino, J.M., Angenent, G.C., and Boutiliera, K. (2017). The BABY BOOM transcription factor activates the LEC1-ABI3-FUS3-LEC2 network to induce somatic embryogenesis. Plant Physiol. 175, 848–857. doi: 10.1104/pp.17.00232.

Ikeuchi, M., Sugimoto, K., and Iwase, A. (2013). Plant callus: Mechanisms of induction and repression. Plant Cell. 25, 3159–3173. doi: 10.1105/tpc.113.116053.

Iwase, A., Mitsuda, N., Koyama, T., Hiratsu, K., Kojima, M., Arai, T., Inoue, Y., Seki, M., Sakakibara, H., Sugimoto, K., et al. (2011). The AP2/ERF transcription factor WIND1 controls cell dedifferentiation in arabidopsis. Curr. Biol. 21, 508–514. doi: 10.1016/j.cub.2011.02.020.

Iwase, A., Mita, K., Nonaka, S., Ikeuchi, M., Koizuka, C., Ohnuma, M., Ezura, H., Imamura, J., and Sugimoto, K. (2015). WIND1-based acquisition of regeneration competency in Arabidopsis and rapeseed. J. Plant Res. 128, 389–397. doi: 10.1007/s10265-015-0714-y.

Kawashima, T., and Goldberg, R.B. (2010). The suspensor: not just suspending the embryo. Trends Plant Sci. 15, 23–30. doi: 10.1016/j.tplants.2009.11.002.

Koi, S., Hisanaga, T., Sato, K., Ishizaki, K., Kohchi, T., Nakajima, K., Shimamura, M., and Yamato, K.T. (2016). An Evolutionarily Conserved Plant RKD Factor Controls Germ Cell Differentiation. Curr. Biol. 26, 1775–1781. doi: 10.1016/j.cub.2016.05.013.

Köszegi, D., Johnston, A.J., Rutten, T., Czihal, A., Altschmied, L., Kumlehn, J., Wüst, S.E.J., Kirioukhova, O., Gheyselinck, J., Grossniklaus, U., et al. (2011). Members of the RKD transcription factor family induce an egg cell-like gene expression program. Plant J. 67, 280–291. doi: 10.1111/j.1365-313X.2011.04592.x.

Lakshmanan, K.K., and Ambegaokar, K.B. (1984). Polyembryony. In Embryology of Angiosperms. 445–474. doi: 10.1007/978-3-642-69302-1_9.

Laux, T., Mayer, K.F.X., Berger, J., and Jürgens, G. (1996). The WUSCHEL gene is required for shoot and floral meristem integrity in Arabidopsis. Development. 122, 87–96.

Liu, Y., Li, X., Zhao, J., Tang, X., Tian, S., Chen, J., Shi, C., Wang, W., and Zhang, L. (2015). Direct evidence that suspensor cells have embryogenic potential that is suppressed by the embryo proper during normal embryogenesis. Proc. Natl. Acad. Sci. USA. 112, 12432–12437. doi: 10.1073/pnas.1508651112.

Llavata-Peris, C., Lokerse, A., Möller, B., De Rybel, B., and Weijers, D. (2013). Imaging of phenotypes, gene expression, and protein localization during embryonic root formation in arabidopsis. Methods Mol. Biol. 959, 137–148. doi: 10.1007/978-1-62703-221-6_8.

Mayer, K.F.X., Schoof, H., Haecker, A., Lenhard, M., Jürgens, G., and Laux, T. (1998). Role of WUSCHEL in regulating stem cell fate in the Arabidopsis shoot meristem. Cell. 95, 805–815. doi: 10.1016/S0092-8674(00)81703-1.

Méndez-Hernández, H.A., Ledezma-Rodríguez, M., Avilez-Montalvo, R.N., Juárez-Gómez, Y.L., Skeete, A., Avilez-Montalvo, J., De-La-Peña, C., andLoyola-Vargas, V.M. (2019). Signaling overview of plant somatic embryogenesis. Front. Plant Sci. 10:77. doi: 10.3389/fpls.2019.00077.

Palovaara, J., de Zeeuw, T., and Weijers, D. (2016). Tissue and Organ Initiation in the Plant Embryo: A First Time for Everything. Annu. Rev. Cell Dev. Biol. 32, 47–75. doi: 10.1146/annurev-cellbio-111315-124929.

Rademacher, E.H., Lokerse, A.S., Schlereth, A., Llavata-Peris, C.I., Bayer, M., Kientz, M., FreireRios, A., Borst, J.W., Lukowitz, W., Jürgens, G., et al. (2012). Different Auxin Response Machineries Control Distinct Cell Fates in the Early Plant Embryo. Dev. Cell. 22, 211–222. doi: 10.1016/j.devcel.2011.10.026.

Radoeva, T., and Weijers, D. (2014). A roadmap to embryo identity in plants. Trends Plant Sci. 19, 709–716. doi: 10.1016/j.tplants.2014.06.009.

Radoeva, T., Hove, C.A.T., Saiga, S., and Weijers, D. (2016). Molecular characterization of arabidopsis GAL4/UAS enhancer trap lines identifies novel cell-type-specific promoters. Plant Physiol. 171, 1169–1181. doi: 10.1104/pp.16.00213.

Radoeva, T., Lokerse, A.S., Llavata-Peris, C.I., Wendrich, J.R., Xiang, D., Liao, C.Y., Vlaar, L., Boekschoten, M., Hooiveld, G., Datla, R., et al. (2019). A robust auxin response network controls embryo and suspensor development through a basic helix loop helix transcriptional module. Plant Cell 31, 52–67. doi: 10.1105/tpc.18.00518.

de Rybel, B.D., van Den Berg, W., Lokerse, A., Liao, C.Y., Van Mourik, H., Möller, B., Peris, C.L., and Weijers, D. (2011). A versatile set of ligation-independent cloning vectors for functional studies in plants. Plant Physiol. 156, 1292–1299. doi: 10.1016/j.tplants.2014.06.009.

Schwartz, B.W., Yeung, E.C., and Meinke, D.W. (1994). Disruption of morphogenesis and transformation of the suspensor in abnormal suspensor mutants of Arabidopsis. Development. 120, 3235–3245.

Snapp, E.L. (2009). Fluorescent proteins: a cell biologist’s user guide. Trends Cell Biol. 19, 649–655. doi: 10.1016/j.tcb.2009.08.002.

Tedeschi, F., Rizzo, P., Rutten, T., Altschmied, L., and Bäumlein, H. (2017). RWP-RK domain-containing transcription factors control cell differentiation during female gametophyte development in Arabidopsis. New Phytol. 213, 1909–1924. doi: 10.1111/nph.14293.

Vernon, D.M., and Meinke, D.W. (1994). Embryogenic Transformation of the Suspensor in twin, a Polyembryonic Mutant of Arabidopsis. Dev. Biol. 165, 566–573. doi: https://doi.org/10.1006/dbio.1994.1276.

Völz, R., Heydlauff, J., Ripper, D., vonLyncker, L., and Groß-Hardt, R. (2013). Ethylene Signaling Is Required for Synergid Degeneration and the Establishment of a Pollen Tube Block. Dev. Cell. 25, 3210–316. doi: 10.1016/j.devcel.2013.04.001.

de Vries, S.C., and Weijers, D. (2017). Plant embryogenesis. Curr. Biol. 27, 870–873. doi: 10.1016/j.cub.2017.05.026.

Waki, T., Hiki, T., Watanabe, R., Hashimoto, T., and Nakajima, K. (2011). The arabidopsis RWP-RK protein RKD4 triggers gene expression and pattern formation in early embryogenesis. Curr. Biol. 21, 1277–1281. doi: 10.1016/j.cub.2011.07.001.

Weijers, D., vanHamburg, J.P., vanRijn, E., Hooykaas, P.J.J., and Offringa, R. (2003). Diphtheria toxin-mediated interregional communication seed development. Plant Physiol. 133, 1882–1892. doi: 10.1104/pp.103.030692.2003.

Weijers, D., Schlereth, A., Ehrismann, J.S., Schwank, G., Kientz, M., and Jürgens, G. (2006). Auxin triggers transient local signaling for cell specification in Arabidopsis embryogenesis. Dev. Cell. 10, 265–270. doi: 10.1016/j.devcel.2005.12.001.

Wendrich, J.R., Möller, B.K., Uddin, B., Radoeva, T., Lokerse, A.S., De Rybel, B., and Weijers, D. (2015). A set of domain-specific markers in the Arabidopsis embryo. Plant Reprod. 28, 153–160. doi: 10.1007/s00497-015-0266-2.

Wickramasuriya, A.M., and Dunwell, J.M. (2015). Global scale transcriptome analysis of Arabidopsis embryogenesis in vitro. BMC Genomics. 16:301. doi: 10.1186/s12864-015-1504-6.

Xu, K., Liu, J., Fan, M., Xin, W., Hu, Y., and Xu, C. (2012). A genome-wide transcriptome profiling reveals the early molecular events during callus initiation in Arabidopsis multiple organs. Genomics. 100, 116–124. doi: 10.1016/j.ygeno.2012.05.013.

Zuo, J., Niu, Q.W., Frugis, G., and Chua, N.H. (2002). The WUSCHEL gene promotes vegetative-to-embryonic transition in Arabidopsis. Plant J. 30, 349–359. doi: 10.1046/j.1365-313X.2002.01289.x.

